# Origin and evolution of Auxin Efflux Carrier family: PIN, PILS, GPR155 and the others

**DOI:** 10.1101/2024.06.26.600818

**Authors:** Stanislav Vosolsobě, Katarina Kurtović, Vojtěch Schmidt, Roman Skokan, Jan Petrášek

## Abstract

The resolution of the 3D structure of PIN proteins and the ability to universally predict protein structures using AlphaFold allowed us to reconsider the classification of Auxin efflux carriers within the BART superfamily, and we propose to merge members of this superfamily, possessing a characteristic fold with a helical crossover, with other families carrying this fold (BASS, NhaA, CPA and others) into a newly proposed superfamily named X-Helices Carriers (XHC). We further demonstrate a profound divergence between PIN and PILS proteins and monophyly of eukaryotic PINs with the animal cholesterol receptor GPR155/LYCHOS. We hypothesise that its signalling capacity is derived from a general feature of PIN proteins to interact with membrane lipids within the core membrane fold. Finally, we discuss the auxin-transport ability of PIN and PILS proteins in the context of the described properties of bacterial homologs that act as organic acid permeases.

**Disclaimer:** This version is a draft and it will be updated during the next few weeks.

**Highlights:**

- PINs are in plants and fragmentary in many algae and protists, but never in fungi
- Eukaryotic PINs are monophyletic together with animal cholesterol receptor GPR155
- PILS are frequently present in protists and in all fungi, but not in animals
- PINs/PILS split is occurred very early in Prokaryota
- AECs belongs to proposed superfamily of transporter with helical crossover

## Introduction

Where did the auxin signalling come from? If we look outside the terrestrial plant, we do not find homologs of the canonical auxin F-box protein family receptor-based signalling pathway, nor do we even find homologs of key genes for auxin synthesis. However, we do find the IAA and its metabolites across the whole green lineage (Schmidt et al., 2024) and an apparent conservation of auxin carrier genes ensuring auxin export from the cytoplasm, namely, proteins from the PIN-FORMED (PIN) and PIN-LIKES (PILS) families (Carrillo‐Carrasco et al., 2023; Vosolsobě et al., 2020). While the former family has been intensively studied for several decades, as its members are among the most important morphogenetic regulators in plants, PILS were discovered only a decade ago (Barbez et al., 2012; Feraru, 2012) and, in contrast to PINs, only provide homeostatic regulation of auxin levels in the cell, not its export from the cell to the environment.

Evolutionary reconstruction of the PIN-FORMED family is complicated by the convergent origin of truncated paralogs (Bennett et al., 2014) that do not contain a regulatory cytoplasmic loop and which are localised on the endoplasmic reticulum membrane, not on the plasma membrane. In initial phylogenetic analyses, these proteins clustered together at the base of the phylogenetic tree with homologs from *Physcomitrium*, for which only ER localization was erroneously detected. This led to the idea that their role in regulating intracellular auxin homeostasis is also ancestral in the PIN family. When the ability to bind to the plasma membrane was demonstrated in both moss PINs and the first characterised algal representative from *Klebsormidium flaccidum* (Skokan et al., 2019), which also possesses the demonstrated ability to transport IAA, it appears as a plausible hypothesis that the export of IAA from the cell may have been an ancestral role of these proteins long before the origin of plants.

Insights from annotation of streptophyte algal genomes indicated that the PIN family is deeply conserved in this algal lineage (Vosolsobě et al., 2020). However, apparent homologs have long been known from bacteria or from *Trichomonas vaginalis* and are collectively summarised as Auxin Efflux Carriers (AEC). More recently, a number of homologues have also been identified in algae outside the Streptophyta lineage (Bogaert et al., 2022). However, the evolutionary relationships within the AEC family have not been further investigated by dedicated phylogenetic analysis. The observed disjointed pattern of homologue distribution across eukaryotes can be explained either by a series of horizontal gene transfers (HGTs) or by parallel gene losses.

Although the PILS family of carriers was identified on the basis of structural similarity to PINs (Barbez et al., 2012), their evolutionary separation took place very long ago at the bacterial level (Bogaert et al., 2022). The whole AEC family is classified in the BART superfamily within the system of Transporter Classification Database (Saier et al., 2021), whose members were originally Bile/Arsenite/Riboflavin Transporters, but currently, bile acid/sodium symporters (BASS) are classified into IT superfamily. The validity of this classification could be recently validated, using AlphaFold prediction, its deduction of the 3D structure in case of PIN proteins is in absolute agreement with conserved NhaA-type fold obtained by cryo-EM (Su et al., 2022; Ung et al., 2022; Yang et al., 2022).

The aim of this paper is to fill the gaps in the description of the evolutionary relationships between eukaryotic and prokaryotic representatives of AECs and the relationship of the AEC family to other members of the BART superfamily. We collected both positive and negative data on the presence of PIN homologs from all available annotated eukaryotic genomes, performed extensive phylogenetic analysis, and analysed the structures predicted by AlphaFold for representative AEC family representatives in the light of recently published structure and mechanism of function of plant PINs. Finally, for the first time, the relationship of PIN, PILS and animal G-protein coupled receptor GPR155 is critically discussed in the light of recent results showing its role in cholesterol sensing in lysosomes (Shin et al., 2022).

## Results and Discussion

### Reclassification of BART superfamily and definition of new XHC superfamily

With the primary goal of finding an outgroup to the AEC family due to the rooting of its phylogenetic tree, we decided to collect representatives of all related transporter families. The existing classification of superfamilies proved to be completely inadequate, as the structure of proteins classified into individual families was extremely variable and did not satisfy our criterion that the presence of an NhaA-type fold with 10 TM helices with the central crossover would be a sign of phylogenetic relationship.

Using natural intelligence, we analysed experimentally-obtained or AlphaFold-computed folds for representative proteins from all classes of membrane transporters listed in TCDB (Saier et al., 2021) and we selected all classes having features of NhaA-type fold. Matching structures were found within BART superfamily - 2-Keto-3-Deoxygluconate Transporters (KdgT, 2.A.10) and Auxin Efflux Carriers (AEC, 2.A.69), in IT superfamily - Bile Acid:Na+ Symporters (BASS, 2.A.28) and Arsenical Resistance-3 transporters (ACR3, 2.A.59), in CPA superfamily - Monovalent Cation:Proton Antiporters (CPA1 & CPA2, 2.A.36 & 2.A.37) and finally within unclassified transporters - NhaA Na+:H+Antiporter (NhaA, 2.A.33) and Na+-dependent Bicarbonate Transporter (SBT, 2.A.83); Fig. 1. On the other hand, Sensor Histidine Kinases (SHK, 9.B.33) and Kinase/Phosphatase/Cyclic-GMP Synthase/Cyclic di-GMP Hydrolases (KPSH, 9.B.34) from BART superfamily and all other members of IT and CPA superfamilies exhibit different fold.

**Fig. 1:**
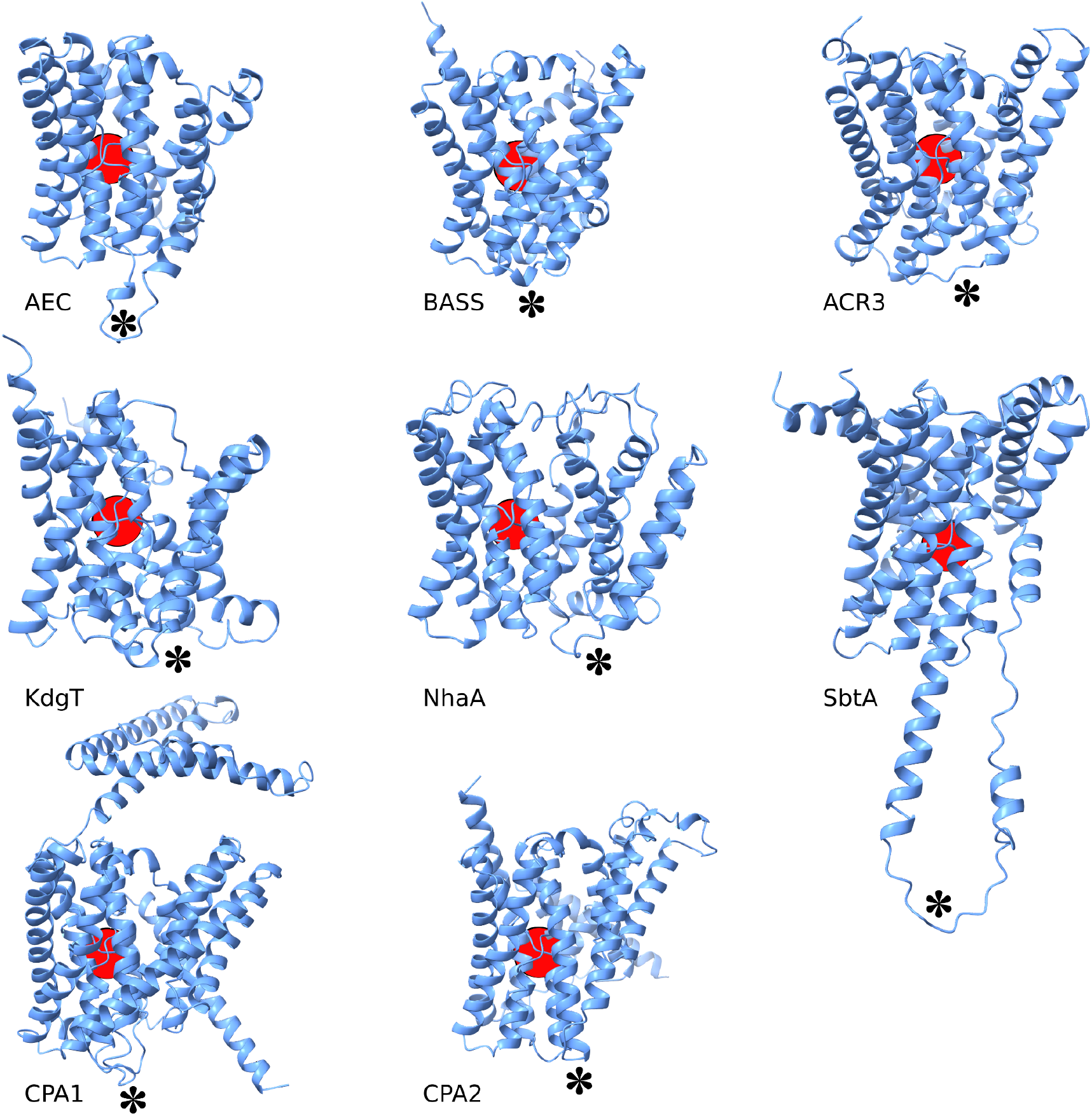
Comparison of core protein structures in proposed XHC superfamily. AlphaFold predicted structures were placed in equal orientation, central helical crossover is shown by red circle, position homologous to intracellular loop between fifth and sixth helices is shown by asterisk. AEC - *Escherichia coli* P0AA50, BASS - *Neisseria meningitidis* Q9K0A9, ACR3 - *Shewanella oneidensis* Q8EJD3, KdgT - *Gardnerella vaginalis* S4H275, NhaA - *Geobacter metallireducens* Q39WP5, SbtA - *Microcystis aeruginosa* P0AA50, CPA1 - *Synechocystis sp*. P73863, CPA2 - *Thermus thermophilus* Q72IM4.

Based on these findings, we propose the definition of the new transmembrane protein superfamily named X-Helices Carriers (XHC) including NhaA, SBT, CPA1, CPA2, ACR3, BASS, AEC and KdgT families, which may have ancestral origin and exhibit a conserved elevator-type of ligand transfer mechanism.

### Phylogenetic relationships within proposed XHC superfamily

In order to uncover the relationship between all transporters bearing central crossover, we used aligned representatives of each TCDB family as a query for HMMER search and collected the full set of their homologs available in the Reference proteomes database (in total, 39,784 sequences with a length at least 100 aa and *E-value* of HMMER search < 1.0E-10).

For phylogenetic analysis, this collection was reduced by USEARCH clustering at the level of 45% identity to get 2,326 centroids which were subjected to three consecutive MAFFT G-INS-1 alignments, with removement of all sequences with incomplete coverage of in transmembrane regions or with longer unique insertions (may be a result of incorrect splicing prediction). Refined dataset were subjected to more precise iterative alignments, when all three available algorithms (global pair, local pair and genaf pair) were used in combination with three different BLOSUM matrices. The alignment produced by the genaf pair method proved to be too artificial, so we used only the outputs of the remaining two methods for selection of parsimony informative positions by two methods with different stringency. Resulted 12 alignments were subjected to IQ-TREE maximum likelihood phylogenetic inference with LG+I+R substitution model and 50% majority-rule consensus tree was derived (Fig. 2A).

**Fig. 2:**
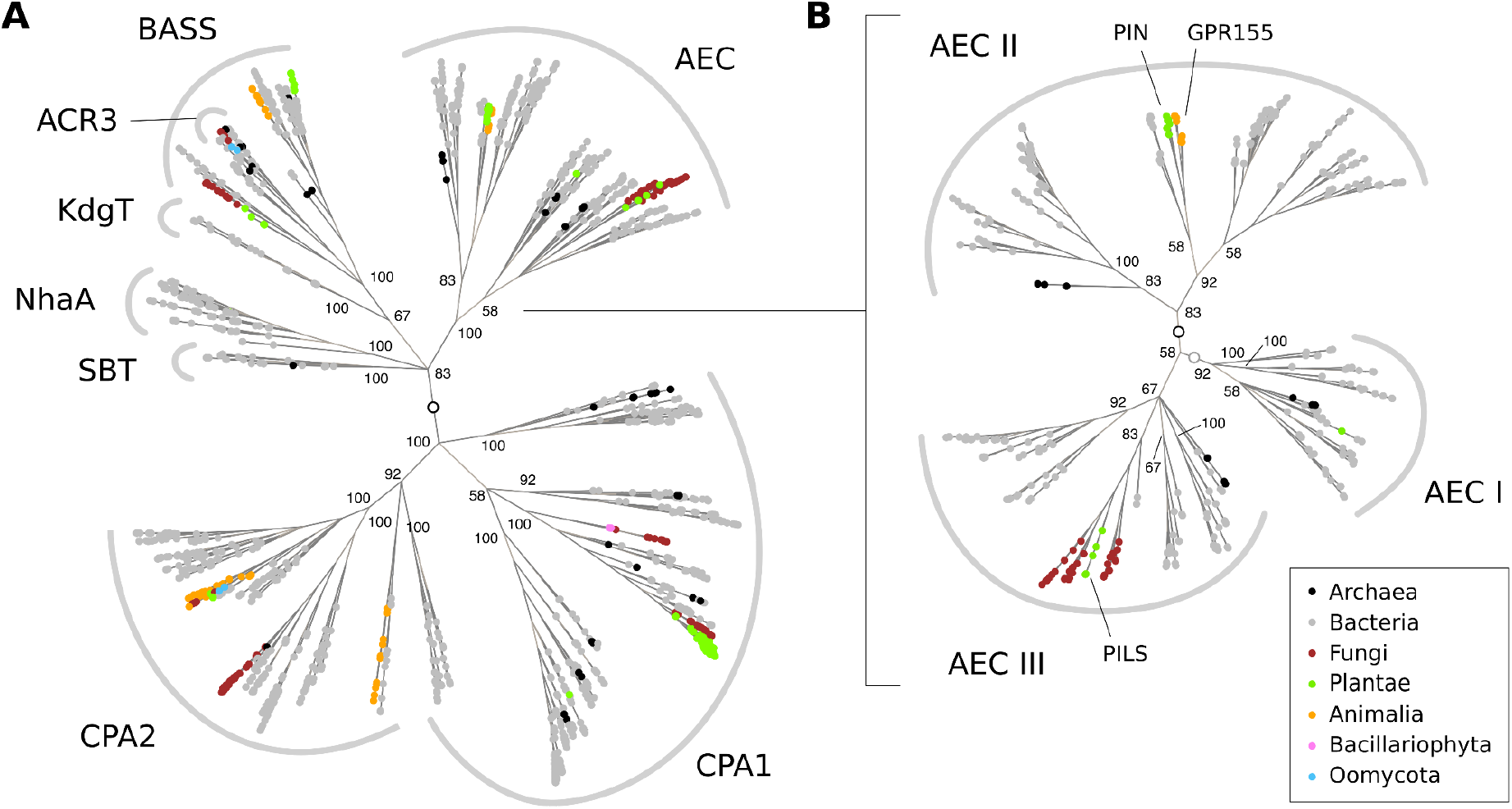
Phylogeny of XHC superfamily. (A) a 50% majority-rule consensus tree derived from individual ML trees based on 12 different alignments computed by G-INS-i or L-INS-i algorithm with three different BLOSUM matrices (BLOSUM30, 45 and 62) and using all positions selected by dynamic determination of gaps threshold or only positions with gappiness < 95 % in Clipkit. Empty circle shows the root determined by mid-point. (B) A subtree representing the AEC family. Dark circle indicates the root supported by 5 of 12 topologies, light circle indicates root supported by 3 topologies. Node labels indicate the percentage of monophyly of the given clade in the consensus tree.

There was a big variability in the topology of clade branching among the XHC trees. Unquestionably, there is no justification for a separate definition of the ACR3 family, as it is an internal group of the BASS family. The KdgT family shows affinity to BASS and the CPA1 and CPA2 families form a monophyletic family, but the affiliation of some lineages to CPA1 or CPA2 is not clear. In contrast to previously published (Masrati et al., 2018), we did not demonstrate affinity between NhaA and CPA2. The AEC line itself is very clearly distinguished from other groups and has no higher affinity to any particular group.

### Rooting of AEC family and its phylogeny

Using the previously derived consensual tree, the position of the root of the AEC family was estimated (Fig. 2B). Despite partial discrepancies in the resolution of the main AEC clades, we can state that the AEC family is formed by three main line, by one solely prokaryotic AEC I, and two with both prokaryotic and eukaryotic representatives, while the first AEC II include whole PIN family and the second, more diversified AEC III with the PILS family, both accompanied by several clades of prokaryotic homologues. The position of the root was most frequently placed in the branch leading to AEC II (5 of 12 topologies), next in the branch to AEC I (3 of 12 topologies). Alternate topologies with the root in the branches leading to PILS, GPR155, AEC III or between two clades of AEC II were obtained in single cases. Support for a monophyly was 92 % for AEC I, 83 % for AEC II and 67 % for AEC III. Thus, we can conclude that the divergence between PIN and PILS is a major gradient within the AEC family.

After estimation of its root, we computed unrooted phylogeny of solely AEC homologs (Fig. 3) to get a better resolution of internal relationships. In the clade AEC II, there is one cluster of sequences with higher affinity to the eukaryotic cluster PIN & GPR155 (AEC IIb). In AEC III, we resolved cluster AEC IIId, which is sister to eukaryotic PILS; the topology of the other main clusters (AEC IIIa-c) is variable and not clearly resolved.

**Fig. 3:**
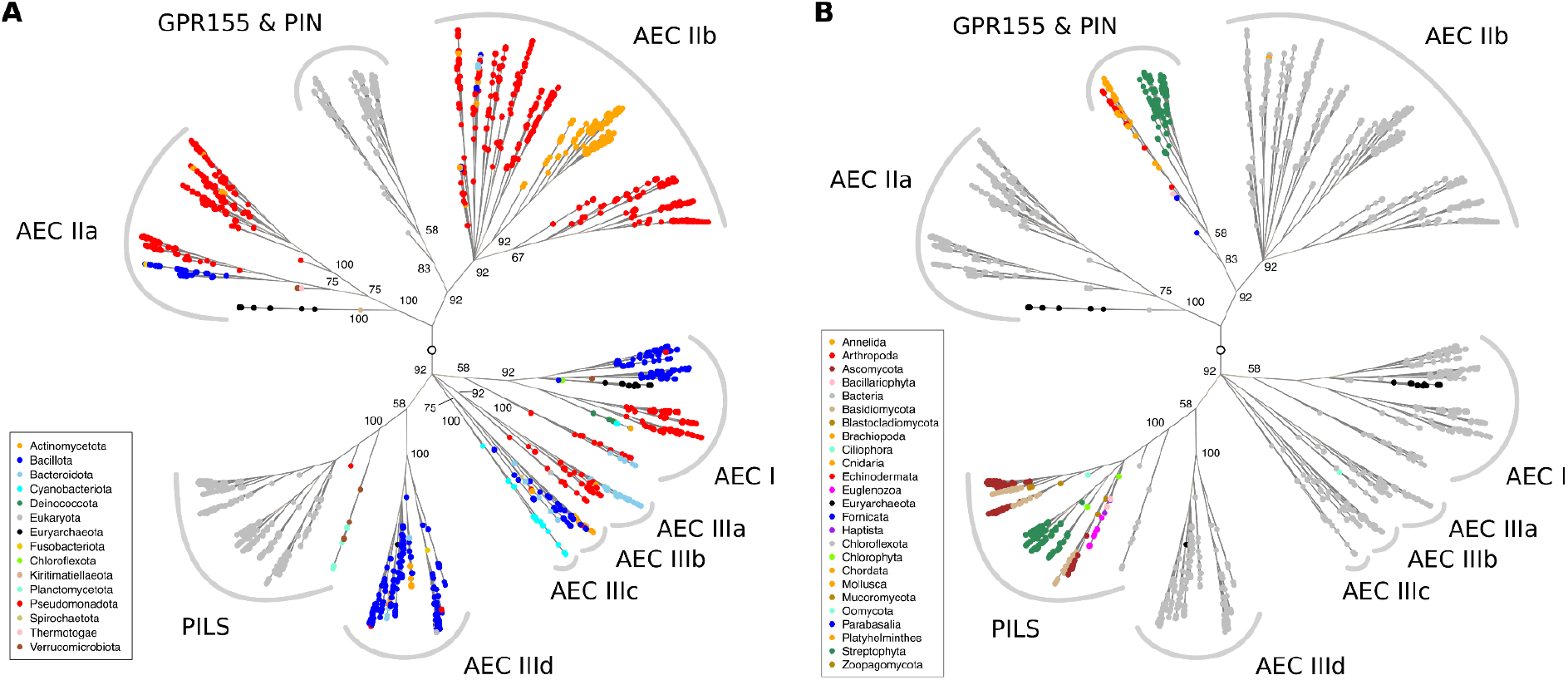
Phylogeny of AEC family. A 50% majority-rule consensus tree derived from individual ML trees based on 12 different alignments computed by G-INS-i or L-INS-i algorithm with three different BLOSUM matrices (BLOSUM30, 45 and 62) and using all positions selected by dynamic determination of gaps threshold or only positions with gappiness < 95 % in Clipkit. Empty circle shows the most probable root position determined previously. Eukaryotic and prokaryotic lineages are colour-coded in (A) and (B) respectively.

### Divergency of PIN and PILS and the ancestral function of AEC

Despite the fact that PIN and PILS proteins were described to play similar roles in auxin transport in plants, all obtained topologies in our phylogenetic analysis clearly show that these are lineages separated deeply at the prokaryotic level (Fig. 3) in agreement with previous studies (Bogaert et al., 2022; Feraru, 2012). Eukaryotic PINs are sister to the clade present mainly in Psudomonadota (form. Proteobacteria). It includes MdcF transporter, which is coded by the operone responsible for malonate utilisation in *Klebsiella pneumoniae* and thought to be a malonate importer (Hoenke et al., 1997), but which is not present in the other species utilising malonate (Chohnan et al., 1999; Stoudenmire et al., 2017), and YfdV, transporter coded by operone ensuring the survival of *Escherichia coli* exposed to oxalic acid stress (Fontenot et al., 2013). However, the characterization of both proteins is poor.Next member of this clade, homolog of Bacillus licheniformis YwkB (Q65DU8), was found by hot spring metagenome sequencing and predicted to be an auxin transporter by ligand docking and MD simulation (Rai et al., 2021). The presence of YwkB in *Paenibacillus polymyxa* was found to be necessary for plant growth-promoting activity in this Rhizobacteria (Da Mota et al., 2008). Despite the ligand similarity, there is no phylogenetic affinity of these proteins to plant PINs.

The group of the closest bacterial homologs of PILS is represented by malate uptake carrier MleP in wine lactobacteria Oenococcus oeni (Labarre et al., 1996), which is driven by proton-motive force in uniport mechanism (Salema et al., 1994) as described in PINs (Ung et al., 2023).

Taken together, bacterial AEC are very poorly characterised and a wide variety of transported substrates can be found among them. Their import function is probably prevalent, however, in the case of rhizobacteria, direct auxin export can also be anticipated. Based on the available knowledge, we can speculate that a general feature of AECs is the ability to transfer organic acid ions, taking advantage of the existing electrical potential across the membrane and the negative charge of the transferred ion. Due to the coupling to the membrane potential only, we can assume that this will be a very fast transport mechanism with kinetics determined only by the availability of the substrate and not by an additional cofactor, such as in the case of a symport/antiport with Na+. As a result, AECs may act as highly robust transporters, for example, in the case of the need for efficient detoxification, both of the external environment (oxalic acid stress in the case of YfdV) and of the internal environment of the cell, in the context of the hypothesis that auxin export is evolved from the need for cell detoxification.

An alternative hypothesis can be postulated: if AEC transporters allow the secretion of an organic acid ion in the direction of increasing acidity, they also change the acid-base balance in the target environment. Suppose that a pH gradient is created in the direction to the extracellular environment through the membrane H+ATPase, then due to the presence of Cl- as the most common counteranion, a strong acid environment is created. This can have undesirable effects, for example excessive protonation or hydrolysis of extracellular matrix components. If a weak organic acid ion is secreted at the same time, buffering of the environment and better control of the cell above the pH of the extracellular environment will occur.

If the AEC transporter is regulated by post-translational modification, it can be activated very flexibly and allow local regulation of the extracellular pH, while the activity of H+ATPase ensuring the formation of proton-motive force can be constant. Indolyl acetic acid (IAA) could primarily be used for this purpose, as it is i) a secondary metabolite that does not interfere with the basic metabolic ones and ii) due to its large molecular weight, it has a limited ability to migrate into the extracellular matrix, which could be used in organisms with a compact cell wall . The relatively high price of IAA as a nitrogen-containing metabolite speaks against this hypothesis, which, however, might not be a major problem in sessile organisms under terrestrial conditions.

### Distribution of AEC family in Eukaryota

The main goal of this study is to analyse individually all available high-quality proteomes for presence or absence of AEC homologs. To fulfil this aim, we performed both BLAST and HMMER search with the representative set of AEC proteins and their HMM profiles respectively. Our initial intention was to search for PIN homologs in all available complete genomes of eukaryotic organisms, however, BLAST searches in unassembled genomic fragments did not yield reliable results, so we restricted our search to 1648 annotated genomes only available in Reference proteomes collection and on PhycoCosm server. We found a very contrasting pattern, when comparing the PIN and PILS family (Fig. 6).

PIN proteins are present in the land plants (Embryophyta), partially in the other Streptophytes, mainly in Zygnematophyceae and Charophyceae and next, in individual representatives of green algae and diatoms, in Metamonads (*Trichomonas, Histomonas*) Choanoflagellates (*Salpingoeca rosetta*), Ichthyosporea (*Sphaeroforma arctica*) or Myzozoa (*Symbiodinium, Vitrella brassicaformis*).

Typically, their distribution is fragmental outside the Streptophyta and presence of PIN homologue is not a synapomorphy of any higher taxonomic group, with one exception, the GPR155, which is generally conserved in animals. In the PILS family, the presence of homologs is conserved among whole plants (including green algae) with only individual exceptions, e. g. in *Chara braunii*. Outside the green lineage, PILS are also present more frequently, but fragmented as well. In contrast to PINs, PILS are non-present in animals, but absolutely universal in fungi, where only one PIN homolog was found in the whole kingdom, namely in Chytridiomycota. Because this homolog is the most similar to PIN of diatoms or green algae, we are forced to speculate that it was laterally transferred from the prey of this algae-feeding fungi (we note that its similarity to algal homologues is quite distant, which excludes the possibility of contamination during sequencing).

Additional high-quality annotated proteomes will be needed to clearly answer the question of the origin of eukaryotic lineages of AEC proteins. At the same time, it is necessary to keep in mind the fact that, especially among protists, a large number of sequenced organisms are representatives of picoplankton, so they may have a highly reduced number of genes and AEC homologues may be missing secondarily.

### Monophyly of GPR155 with PIN and its origin

Our initial hypothesis was that PIN and GPR155 are independent eukaryotic lines acquired from the prokaryotic AEC pool. Nevertheless, the monophyly of the clade consist of streptophyte PINs, animal GPR155 and several PIN homologs from the eukaryotes, namely other *Fistulifera solaris* (Bacillariophyta), *Trichomonas foetus* (Parabasalia) and *Spironucleus salmonicida* (Fornicata) was not disrupted in 11 from 12 generated topologies and the ultra-fast bootstrap support of this clade varied between 60-80 %. Thus, we are forced to conclude that the whole clade has a unique origin. With respect to fragmentary distribution of PIN homologs among Eukaryota and limited number of known representatives in protozoan taxa, we are unable to formulate a plausible hypothesis for the origin of the animal GPR155 line, but ancestral presence in single eukaryotic group and next the lateral gene transfer between eukaryotic lineages is more probable than hypothesis of universal PIN presence in LECA followed by multiple gene losses.

The role of GPR155 has been unknown for a long time, until its characterisation of the cholesterol receptor in lysosomes connected to mTORC1 signalling cascade (Shin et al., 2022). Cholesterol was found to interact with helix H1 of permease core in GPR155, which is in good agreement with the described binding pocket for membrane lipids in PIN8 (Ung et al., 2022). We speculate that in general, lipids and/or sterols are cofactors necessary for a proper functioning of PIN proteins sensu lato, whose ancestral function was to ensure the flow of organic acids across the membranes. This could be also the case in the ancestor of Metazoa, where PIN homolog could be localised on the membrane of lysosome and regulate its acidity, but subsequently, after its fusion to G-protein domain (Fig. 5E,F), its general lipid-binding ability was co-opted in a new biological role.

**Fig. 5:**
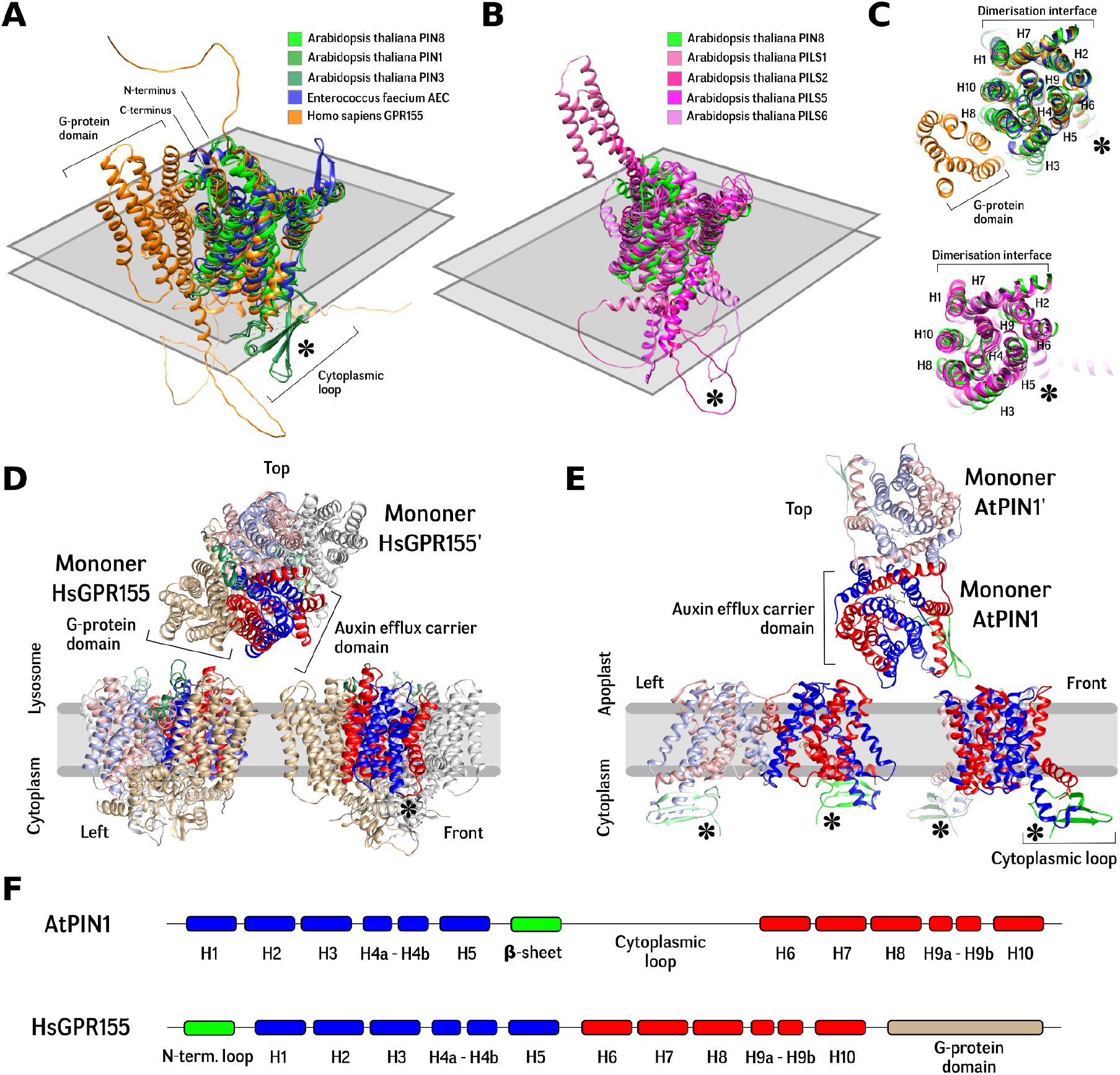
Comparison of structures of AEC proteins. (A) Bacterial AEC proteins (blue) are formed only by 2×5 TM helices. Characteristic feature of plant PIN proteins (green) is a long cytoplasmic loop starting with the β-sheet domain. Animal GPR155 (yellow) are loopless, but lumenal N-terminal extension and big C-terminal membrane G-protein domain is present. (B) Plant PILS are structurally more variable than PIN proteins, cytoplasmic loop is short, but N- or C-terminal extensions are present. (C) Mid-plane cross-section of previous structural alignments. Dimerisation of HsGPR155 (D) and AtPIN1 (E) predicted by AlphaFold and cryo-EM (Yang et al., 2022) respectively. GPR155 has the same dimerisation capability as plant PIN proteins, G-protein domains are located on the sides of the dimer. (F) Schema of domain organisation in AtPIN1 and HsGPR155.

**Fig. 6:**
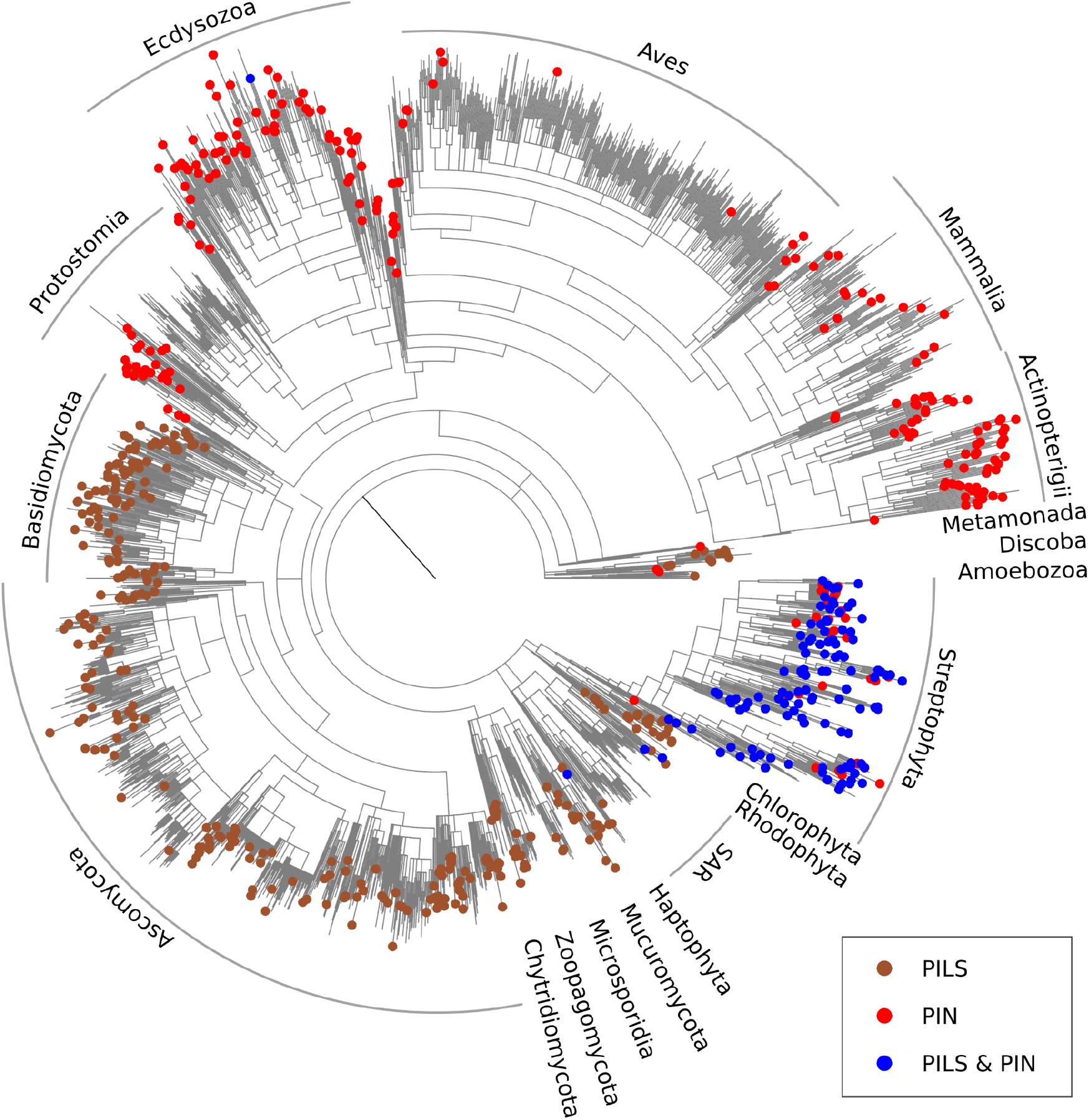
Distribution of PIN and PILS proteins among eukaryotic lineages. NCBI Taxonomy common tree of all taxa with available Reference Proteomes (June 2024) with mapped presence of PIN, PILS or both proteins. The absence of homologs in Aves is given by the outdated Reference Proteomes database (v. 2021) used by the HMMER server.

## Conclusions

Although the last few years have brought a revolution in the knowledge of the mechanism of function of PIN proteins and allowed their structural comparison with other families of membrane transporters, fundamental questions regarding their evolutionary origin remain open. In particular, there is a need to unravel these ambiguities i) what is the role of IAA and related metabolites outside land plants, if it is a waste product, or even a toxic product, or a substance with signalling significance; ii) whether non-plant PIN homologues transport IAA, this ability has so far only been demonstrated in Streptophytes, but it would be important to investigate the properties of the proteins across protists; iii) what is the overall variety of prokaryotic AEC substrates, whether they function as exporters or importers, and whether IAA appears among the substrates in a context other than that of rhizobacteria. Unfortunately, it is necessary to state that these questions will not be answered by the current “genomic era” of EVO-DEVO biology, but it will be necessary to delve deeper into the study of the physiology of non-model organisms, for which their “domestication” in laboratory conditions is often a stumbling block.

## Methods

### Phylogeny of XHT superfamily

The AlphaFold predicted structures of representatives of all TCDB families (Saier et al., 2021) were visually checked for presence of helical crossover and all members of families with similarity to NhaA fold were aligned by online MAFFT with the G-INS-i algorithm (Katoh et al., 2019) and resulting alignments were individually subjected to HMMsearch (Finn et al., 2015) against Reference Proteomes database (online parameters -E 1 --domE 1 --incE 0.01 --incdomE 0.03 --seqdb uniprotrefprot). The full set of unique sequences were fast clustered by USEARCH (Edgar, 2010) at the level of 45 % identity and centroids representing clusters with at least 5 sequences were subjected to online G-INS-1 MAFFT alignment, which were used for manual removement of incomplete or aberrant sequences. Phylogenetically informative positions were selected by ClipKIT (Steenwyk et al., 2020) and used for computation of initial phylogenetic tree by IQ-TREE2 (Nguyen et al., 2015) with UFBoot (Hoang et al., 2018). The LG substitution model was proven to be optimal by ModelFinder (Kalyaanamoorthy et al., 2017). Option ‘-bnni’ was used to reduce the risk of overestimating branch support with UFBoot due to severe model violations. Initial tree was used for the next refinement of our dataset, where the sequences with branches longer than 2 were removed. Final dataset were realigned by more precise iterative algorithms G-INS-i, L-INS-i and E-INS-i by online MAFFT with 2 cycles iterative refinements and with combination of three different scoring matrices, BLOSUM30, BLOSUM45 and BLOSUM62. Two different methods of alignment positions selection were applied on each alignment before ML tree estimation, default selection by ClipKIT as a first, and subsequently, manual subselection of only core protein domains as a second. Final phylogeny trees were computed on all prepared datasets by IQ-TREE2, where only the LG set of substitution models was tested to reduce the computational time. Trees based on G-INS-i and L-INS-i were used for 50% majority rule consensus tree estimation by ape package (Paradis and Schliep, 2019) in RStudio (RStudio Team, 2020).

### Phylogeny of AEC family

The reconstruction of the AEC family was performed in the same way as for the whole XHT superfamily, only the subset of AEC homologs from the primary dataset was used and the clustering was done at the level of 70% similarity with removement of singletons. Majority rule consensus tree was used for selection of representative sequences for each defined clade and HMM profiles were built for classification of the other sequences.

### Exploring the distribution of AEC homologs across Eukaryota

All available Reference proteomes and PhycoCosm assemblies (Grigoriev et al., 2021) were examined individually by BLAST search or HMM scan using a representative set of AEC homologs and HMM profiles respectively. Computation was done on the METACENTRUM computer cluster. Sequences were classified into individual clades of AEC based on best hit in reverse BLAST or highest score in HMMscan. Results were mapped onto a common taxonomy tree built on the NCBI taxonomy server. The ape package (Paradis and Schliep, 2019) in RStudio (RStudio Team, 2020) was used for visualisation of results.

### Visualisation of similarity between AEC proteins

AlphaFold F1 v4 models for all BART representatives (or their closest homologs in the absence of respective model) were subjected to multiple protein structure alignment using mTM-align (Dong et al., 2018) and visualised in UCSF ChimeraX (Meng et al., 2023).

## Acknowledgement

This work was supported by Czech Science Foundation project no. 20-13587S. Computational resources were provided by the e-INFRA CZ project (ID:90254), supported by the Ministry of Education, Youth and Sports of the Czech Republic.

